# Candling analysis of egg development in an endangered bird species Crested ibis (*Nipponia nippon*)

**DOI:** 10.1101/2025.11.17.688759

**Authors:** Yuansi He, Xuebo Xi, Siyi Zeng, Hua Huang, Xiangjiang Zhan, Daiping Wang

## Abstract

Identifying key factors influencing the survival of animals, particularly rare and endangered species is crucial to biodiversity conservation. In birds, hatching failure is pronounced in endangered species. Accurate assessment of egg development and the ability to distinguish non-viable eggs are essential prerequisites for identifying the causes of hatching failure and applying appropriate conservation practices. The Crested ibis (*Nipponia nippon*), a flagship endangered species, has a long history of captive breeding, which has contributed to population recovery. However, little is known about the specific process and characteristics of its egg development. In this study, we provided the first comprehensive description of normal egg development in the Crested ibis, including both the changes observed in unfertilized eggs during incubation and embryonic development via candling (n = 106 eggs, with 1,423 candling images), a commonly used non-invasive method for assessing egg fertility and embryonic status, and offered a practical reference for assessing fertilization status and embryo viability. In addition, we estimated the timing of embryo mortality and found that most deaths occurred during mid-incubation (Day 7∼15) (n = 12, 60.0%) and shortly before hatching (Day 23∼29) (n = 7, 35.0%), highlighting these critical periods that require particular attention. This species-specific documentation of egg development provides a valuable reference for accurately assessing embryonic progress, evaluating environmental effects on survival, and guiding adaptive management, benefiting both captive breeding programs and field conservation efforts.

## Introduction

Biodiversity loss has become an urgent global concern, calling for effective conservation actions, particularly for rare and endangered species. Birds represent the most diverse group of vertebrates, yet approximately 11% of avian species assessed by the IUCN are currently threatened with extinction (IUCN 2025). Identifying the key factors influencing their developing effective strategies for survival are therefore of paramount importance. Egg development is crucial for survival in birds. Both in field studies and captive breeding programs, researchers have frequently observed that hatching failure is common, especially among endangered species that have experienced a genetic bottleneck and inbreeding (Koenig 1982, Briskie & Mackintosh 2004, Marshall *et al*. 2023). Hatching failure generally results from either infertility or embryonic death, which can be attributed to intrinsic factors such as parental gametes quality (Donoghue 1999, Assersohn *et al*. 2021), nutritional imbalances in parents (Wilson 1997) and congenital diseases caused by inbreeding (Fu *et al*. 2019), or extrinsic influences including inappropriate temperature and humidity, bacterial infection, and mechanical shock (Ori 2011). It is essential to timely distinguish viable from non-viable eggs and further assess normal embryonic development. In doing so, we can determine the age of eggs, predict hatch date, identify whether hatching failure stems from fertility or embryo mortality, remove abnormal eggs before contamination occurs, pinpointcritical stages of incubation, evaluate environmental impacts on development, and guide appropriate actions to improve breeding success. A comprehensive understanding of egg development is thus one of the prerequisites for diagnosing the causes of hatching failure and improving conservation outcomes for endangered species.

As the most diverse vertebrate group, birds exhibit substantial interspecific variation in egg size, incubation duration, hatchling mass, and developmental maturity (altricial vs. precocial) (Starck & Ricklefs 1998, Deeming 2007, Cooney *et al*. 2020). Correspondingly, species exhibit great variation in embryonic development as well (Joseph Carl Daniel 1957, Blom & Lilja 2005). Species-specific descriptions of embryonic development are essential to guide further research and improve species management. To date, complete or near-complete egg (embryonic) developmental process has been documented in many avian species using candling, breakout examinationand molecular techniques. For example, detailed anatomical studies have been conducted in species of commercial or research importance, such as like Chicken (*Gallus gallus domesticus*) (Hamburger & Hamilton 1951), Pigeon (*Columba livia*) (Łukasiewicz 2014), Japanese quail (*Coturnix coturnix japonica*) (Ainsworth *et al*. 2010), Guinea fowl (*Numida meleagris*) (Araújo *et al*. 2019), Peking duck (*Anas platyrhynchos domestica*) (Trela *et al*. 2025), and Turkey (*Meleagris gallopavo*) (Mun & Kosin 1960). Also, studies have been performed on representative non-domestic taxa with significant evolutionary or ecological importance, such as Emu (*Dromaius novaehollandiae*) (Nagai *et al*. 2011) and ostrich (*Struthio camelus*) (Deeming 1995), whose embryonic traits can provide insights into the evolution and conservation of developmental characteristics across birds. In addition, species such as American kestrel (*Falco sparverius*), a common North American raptor which is frequently used in experimental studies as a model raptor specie (Bird *et al*. 1984, Pisenti *et al*. 2024), and Zebra finch (*Taeniopygia guttata*), another famous model passerine species that has been widely used in avian researches have been carefully studied (Hemmings & Birkhead 2016, Pei *et al*. 2020), along with other taxa including Blue tit (*Cyanistes caeruleus*) (Hemmings & Birkhead 2016), Great tit (*Parus major*) (Hemmings & Birkhead 2016), Pheasant (*Phasianus colchicus*) (Fant 1957), Society finch (*Lonchura striata domestica*) (Yamasaki & Tonosaki 1988), Mallard (*Anas platyrhynchos*) (Caldwell & Snart 1974), Adélie penguin (*Pygoscelis adeliae*) (Herbert 1967) and Bobwhite quail (*Colinus virginianus*) (Hendrickx & Hanzlik 1965). Nevertheless, most documented species belong to species with commercial importance or laboratory model species. Taxonomically, they belong to Passeriformes, Galliformes, or Anseriformes. In contrast, little is known about egg or embryonic development in endangered species or other taxa, especially flagship species that experienced a serious bottleneck. In fact, investigations on embryonic development in these endangered species are of both evolutionary and conservation importance.

Crested ibis (*Nipponia nippon*) is an iconic and flagship endangered species (Pelecaniformes) that has recovered from an extremely small population(Liu 1981, Li 2023). This species was historically widespread across Northeast Asia (BirdLife-International 2001, Li *et al*. 2014), while in the 20^th^ century, its population declined significantly as a result of habitat destruction and human activity (e.g. illegal hunting, overuse of fertilizer and pesticide) (Archibald *et al*. 1980, Ding 2004, Li *et al*. 2009, Feng *et al*. 2019). It was once presumed extinct after populations in Russia, Korea and Japan consecutively died off (Archibald & Lantis 1979, Anderson 1984). Until 1981, a small wild population of seven individuals, including two breeding pairs and three chicks belonging to one of the couples, was rediscovered in Yangxian County of Shaanxi Province, China (Liu 1981). Since then, extensive conservation efforts have been implemented to save this endangered species, and so far, the global population of wild and captive Crested ibis has been thought to exceed 10,000 (Li 2023). Nonetheless, as a species recovered from an extremely small population, it still suffers from inbreeding depression and low genetic diversity owing to the limited number of founders, which are already believed to contribute to reduced embryo survival and individual fitness (Fu *et al*. 2019, Zheng *et al*. 2024). To determine how factors such as inbreeding affect the Crested ibis and to develop appropriate conservation measures, it is first essential to accurately identify the fertilization status of eggs and the developmental condition of embryos. Despite remarkable progress in population recovery, including long-term successful captive breeding, detailed knowledge of egg development in this species remains limited. Earlier studies on Crested ibis reproduction mainly focused on population growth and husbandry improvements, providing quantitative data such as clutch size, fertilization rate, and hatching rate (Xi *et al*. 2001, Huang *et al*. 2016) or practical information on incubation and chick care (Liu *et al*. 1999, Ding 2004, Huang *et al*. 2006). Aside from one report that summarized embryonic mortality rates across developmental stages and included a few photographs of dead embryos (Yang *et al*. 2020), investigation details of egg and embryonic development have been hardly described in the literature.

Candling is a long-established, non-invasive method for assessing egg fertility and embryonic status in both captive and wild populations (Westerskov 1950, Weller 1956, Mauldin 2002, Ernst *et al*. 2004). It involves using a flashlight or specialized device with a cool light source held against the blunt end of the egg (Hall *et al*. 2023). By examining blood vessel patterns, the silhouette of the embryo, the size of air cell, and changes in opacity, researchers can monitor embryonic development. Unfertilized or dead embryos exhibit distinct visual features, enabling early identification and removal of non-viable eggs (Deeming 1995, Mauldin 2002, Ernst *et al*. 2004). Proper candling has been shown to have minimal effect on hatching success (Reis & Soares 1993). Negative impacts generally result from poor handling practices, such as excessive exposure to room temperature, the use of non-specialized heated light sources, or mechanical shock. When performed correctly, candling is a reliable and safe diagnostic tool during incubation. Although its effectiveness decreases in species with thick-shelled or darkly pigmented eggs (Westerskov 1950), as a non-invasive and convenient identification method, candling remains highly valuable for conservation programs, especially in rare species like the Crested ibis that have stringent requirements on species protection.

In this study, we provided the first comprehensive description and assessment of egg development, which included both the changes observed in unfertilized eggs and embryonic development in the Crested ibis based on the candling images collected throughout the entire incubation period. By analyzing extensive images of eggs with different fates, we aimed to 1) describe and illustrate the details of development stages in normal eggs; 2) identify key diagnostic features that distinguishing non-viable eggs, including infertile eggs and dead embryos, from viable eggs; and 3) estimate the death times of embryos to identify critical periods that require greater attention in breeding programs, based on the information of embryos with accurately recorded death times and characteristic differences observed in candling images. Species-specific documentation of egg development is necessary not only for determining whether hatching failure results from infertility or embryonic death, but also for estimating the timing of embryo loss, evaluating environmental effects on survival, and guiding adaptive management. For endangered species such as the Crested ibis, which had experienced a severe genetic bottleneck, comprehensive knowledge and accurate measurement of fertilization and embryo mortality are the prerequisites for diagnosing the causes of hatching failure, quantifying inbreeding depression, and ultimately improving reproductive outcomes. Such information is valuable for both captive breeding and field conservation efforts. Moreover, it plays an essential role in guiding further research and can provide insights into the evolution and conservation of developmental characteristics across birds.

## Materials and Methods

### Study species

Crested ibis is a medium-sized wading bird characterized by red facial skin and legs, and predominantly white plumage with orange-cinnamon tones on the tail, abdomen and flight feathers. Individuals typically reach sexual maturity at around 3 years old (ranging from 2 to 4 years) (Yu *et al*. 2010). The breeding season generally extends from late February to late June, with mating and nest-building occurring from late February to early March, followed by egg-laying from mid-March to early April (Li & Huang 1986, Shi & Yu 1989, Wang & Shi 1999). The incubation period lasts about 28 days, and it usually takes 30∼40 h for the chick to fully hatch out after breaking the egg (Shi *et al*. 1999, Zhai *et al*. 1999, Huang *et al*. 2016). The breeding pair usually lay one clutch per year, but may produce a replacement clutch if the first is lost early in the season. Each clutch contains 2-4 eggs, laid at one- or two-day intervals (Shi *et al*. 1999). While in captivity, egg production can exceed natural levels—sometimes reaching more than ten—due to egg collection, which stimulates additional laying. The Crested ibis is altricial. At the time of hatching, its eyes have not yet opened, with sparse hair covering its body, and its feet are weak. They need to be raised by adult birds until they are 40 to 45 days old before they can leave the nest (Shi *et al*. 1999, Zhai *et al*. 1999)

### Egg collection and incubation

Eggs were collected from the captive Crested ibis population during the 2025 breeding season at the Crested Ibis Breeding Center at Dongzhai National Nature Reserve, Henan Province, China. All eggs (n = 106) were initially incubated naturally by their parents for average 3.7 days (range: 0 to 23) before being transferred to artificial incubation by trained staffs at the breeding center. The average duration of natural incubation before the eggs were transferred to the artificial incubator was 3.7 days. Among these, 85 eggs (80.2%) were moved to artificial incubation between Day 0 and Day 5 of incubation, 8 eggs (7.5%) between Day 6 and Day 11, 9 eggs (8.5%) between Day 12 and Day 17, and 4 eggs (3.8%) between Day 18 and Day 23. Prior to incubation, each egg was disinfected by immersion in warm water containing potassium permanganate and gently dried. Artificial incubation was carried out in hatching incubators (P-008A-TOKI52, Japen; Rcom Pro 20, Korea) at 37.5∼37.7°C and 55%∼65% relative humidity. To simulate natural cooling behavior, eggs were removed from the incubator every 3 hours between 8:00 to 19:00, cooled for 3 minutes, and then returned.

### Egg candling

Eggs were candled daily using a dedicated egg candler (Zhenye (Huizhou) Industrial Co., Ltd, Huizhou, China), which does not generate heat during the examination, thereby minimizing potential harm to the embryos. Candling was conducted in darkness from both the sharp and blunt end of each egg and finished in less than 3 minutes. Eggs were promptly returned to the incubators after images were taken using an iPhone 15 Pro (Apple Inc., USA). After Day 25 of incubation, when hatching was imminent, continuous candling was discontinued for most eggs to avoid disturbing the hatching process, except for a few infertile eggs and those showing no signs of pipping. During the whole incubation period, we carried out following practices: 1) eggs that emitted a foul odor due to decomposition or showed no signs of embryonic development after 7 days were opened to assess fertility; 2) unhatched eggs remaining after the expected hatching date, or eggs in which the embryo appeared non-viable, were dissected to determine developmental status; 3) for embryos that died during incubation without a precisely known time of death, the approximate timing was inferred from embryo size at necropsy combined with candling records obtained during the incubation period.

### Picture processing

A total of 1,423 images of 106 eggs were taken during the breeding seasons (Table 1). Images were taken throughout the entire incubation process, ensuring the comprehensive monitoring of egg development (Table 2). All the images were cropped and arranged using Adobe Photoshop (Adobe Inc., USA) and Adobe Illustrator (Adobe Inc., USA). Since no scale bars were included during photography, image proportions were calibrated using objects of known size present in the images (e.g., operator’s hands, sample tubes).

**Table 1.**
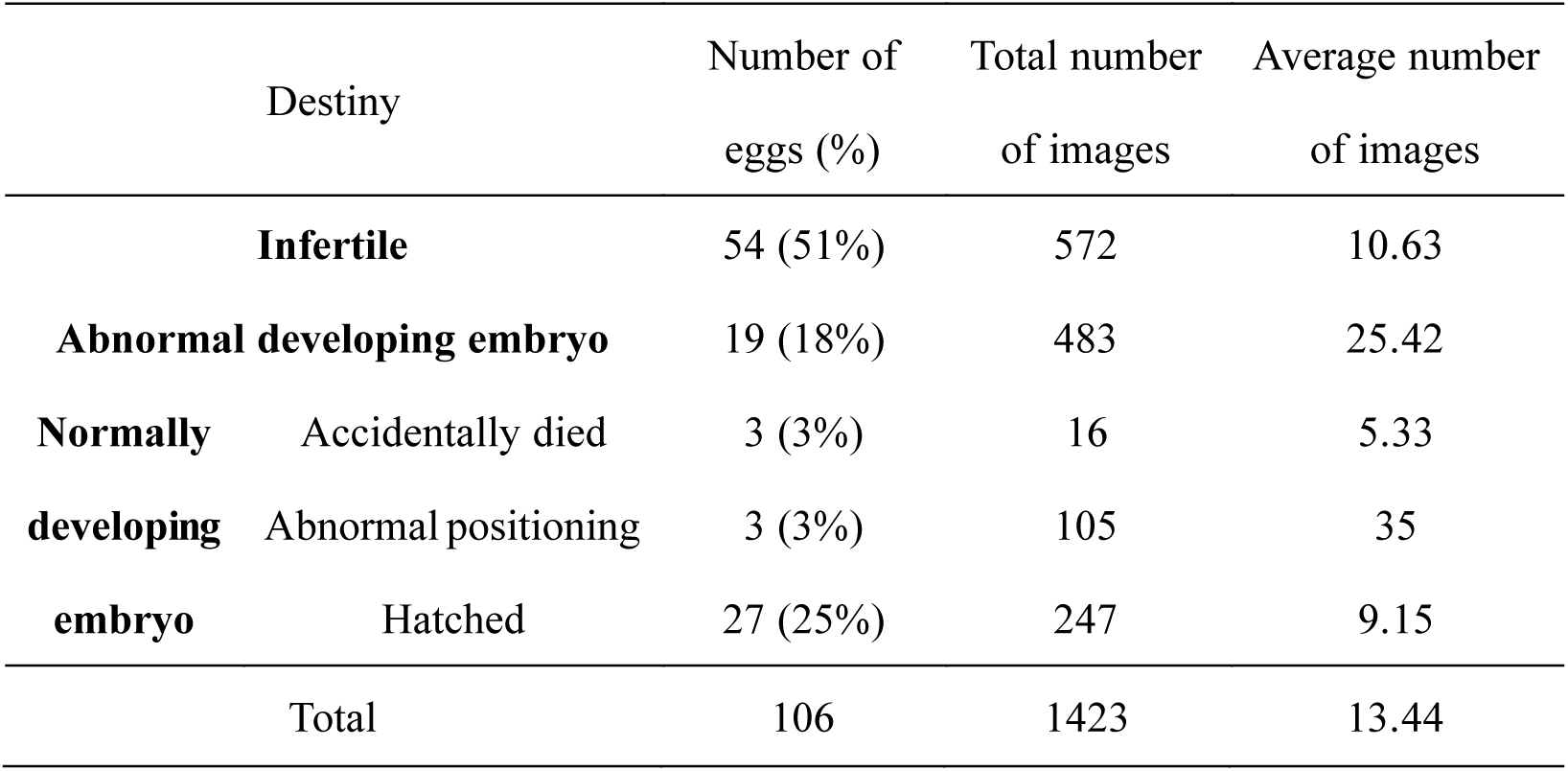
Summary of egg classification, total number of images and average number of images during incubation.

**Table 2.**
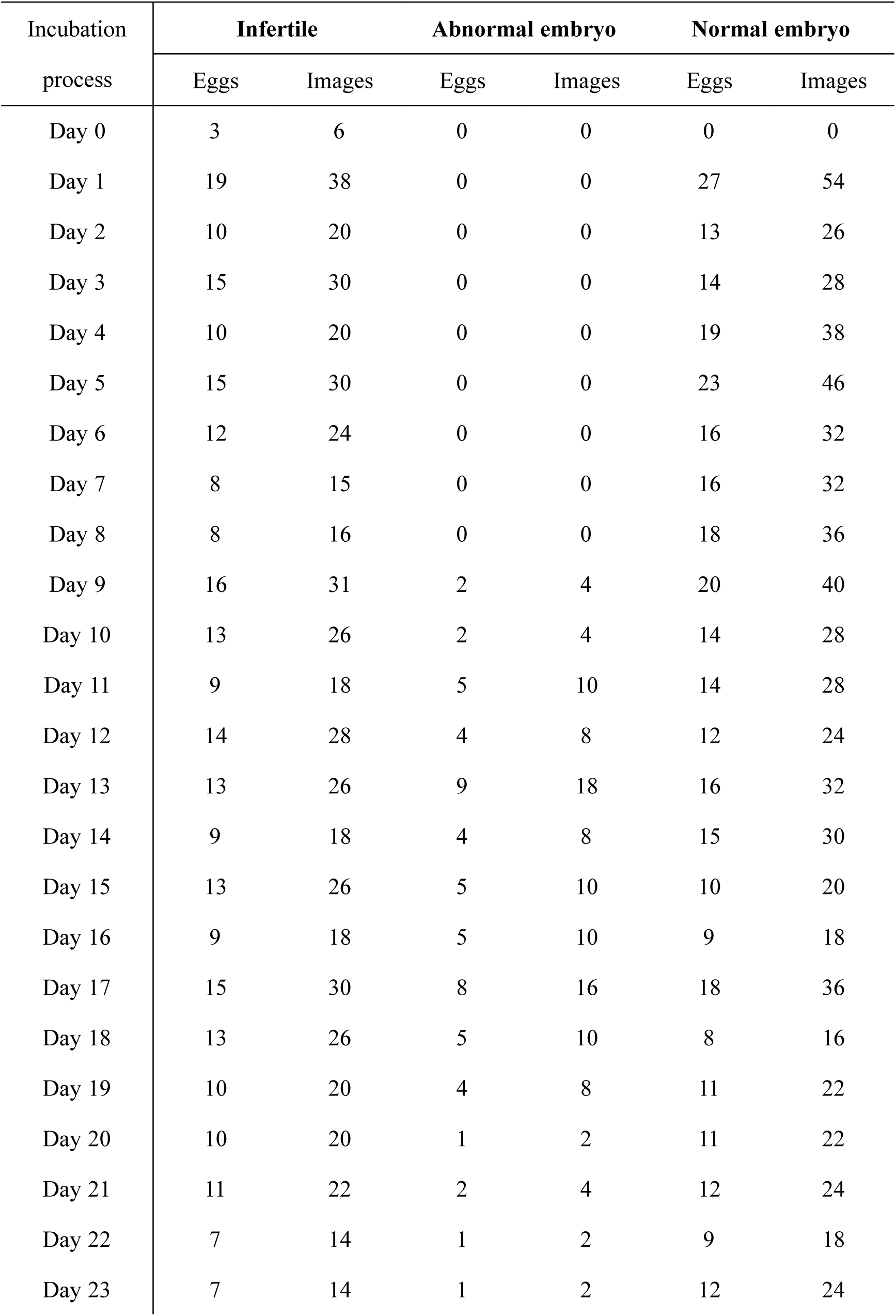
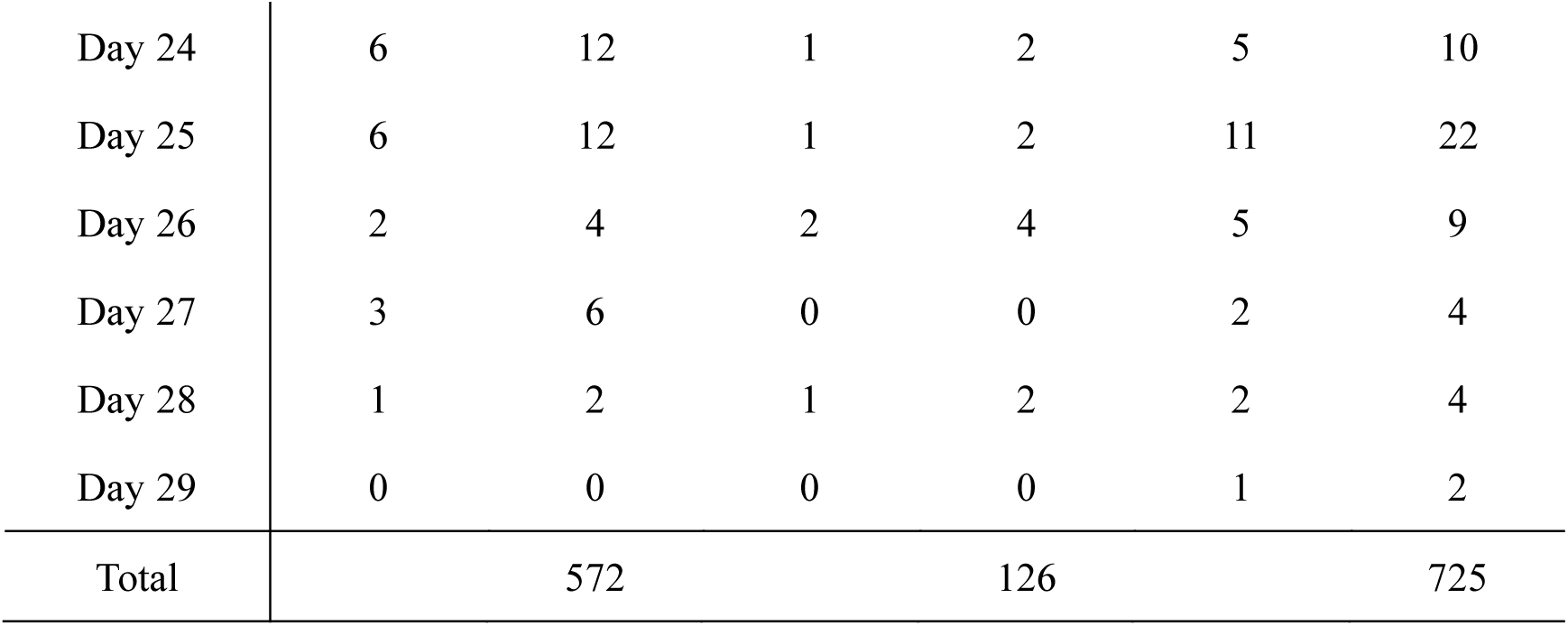
Summary of the number of eggs and images recorded throughout the incubation. Day 0 referred to the day when the egg was laid. For eggs in which the embryos eventually died, images taken before any visible signs of abnormality appeared were classified as **normal embryo**.

The images were first classified into different categories based on the final incubation outcomes and the developmental conditions (i.e. infertile, dead embryo, successfully hatched) determined by break-out examination. To describe and illustrate the developmental details at different stages in normal eggs, we compiled all the candling images of eggs that developed normally (including hatched eggs and those that failed due to accidental breakage or abnormal positioning) to summarize their changing characteristics throughout incubation, and selected representative images for annotation and explanation. Subsequently, to address the key question of how to identify non-viable eggs, including infertile eggs and dead embryos, we compared candling images of normally developing eggs and abnormal eggs (unfertilized or containing dead embryos) at corresponding stages, summarized the key distinguishing features, and chose representative images to illustrate and describe these characteristics. Finally, to determine the timing of embryo death, we first estimated the approximate time of death based on the appearance of abnormalities in the candling images. A more precise estimation was then obtained by comparing the morphological characteristics of the dead embryos after break-out examination with those of four embryos whose death times were clearly recorded due to accidental events. The distribution of death events was subsequently analyzed to identify the critical periods of embryonic development in the Crested ibis. Graphical presentations of the statistical results were produced using the **ggplot2** (Wickham 2016) package in R 4.5.1 (R Core Team 2022).

## Results

### General summary

A total of 106 eggs were collected during the breeding season, of which 52 (49%) were fertilized and 27 successfully hatched (25%). Among the fertilized eggs that failed to hatch (n = 25), three were accidentally broken during handling, and three failed to pip the eggshell because of abnormal fetal positioning, resulting in death from asphyxiation. In total, 1,423 images were taken during the incubation period, including 483 images documenting the developmental process of normally developing embryos (some of which later failed due to accidents or abnormal positioning), 368 documenting embryos that died during development, and 574 documenting the incubation process of infertile eggs (Table 1). In addition, 24 images of dead embryos were taken after egg opening (including one naturally incubated egg that broke because the nest collapsed), allowing partial insight into embryonic morphology at different developmental stages in the Crested ibis.

### a) describing different development stages in normal eggs

The text below describes the candling images of normally developing embryos during the incubation process, which can offer references for assessing fertilization status, estimating embryonic age, and evaluating embryonic viability. For each egg, the images on the top were taken when candling from the sharp end, and those on the bottom were taken when candling from the blunt end.

**Day 1∼3:** Fertilized (n = 31 eggs, 108 pictures; Figure 1a) and unfertilized(n = 26 eggs, 88 pictures; Figure 1b) eggs appear indistinguishable regardless of the candling end, with complete transparency and only a vague yolk shadow without clear boundaries.

**Figure 1.**
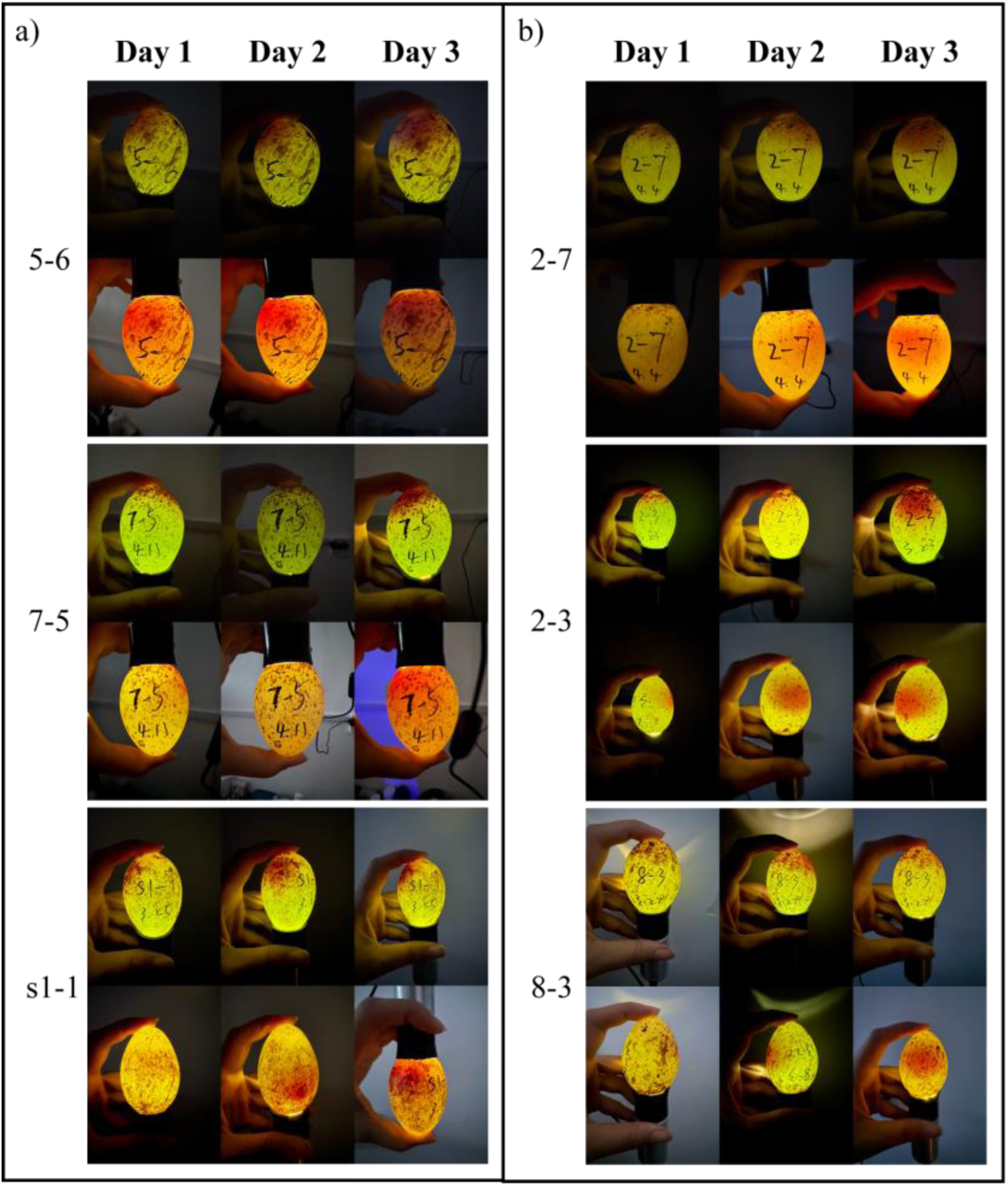
Representative candling images of fertile (a) and infertile (b) eggs during Day 1 to Day 3 of development. For each egg (5-6, 7-5, s1-1, 2-7, 2-3, 8-3), the images on the top were taken when candling from the sharp end, and those on the bottom were taken when candling from the blunt end.

**Day 4∼6:** A distinct shadow becomes visible at the blunt end when candled from the sharp end (Figure 2a-2c, white boxes), indicating the onset of embryonic development. From the blunt end, the shadow at the sharp end appears as a continuous area rather than a floating sphere (Figure 2a, white braces). The shadow is typically darker at the bottom (Figure 2b-2c, dark blue braces), lighter at the top (Figure 2b-2c, pink braces), and a transparent region corresponding to the air cell is visible at the blunt end (Figure 2c, white arrows).

**Figure 2.**
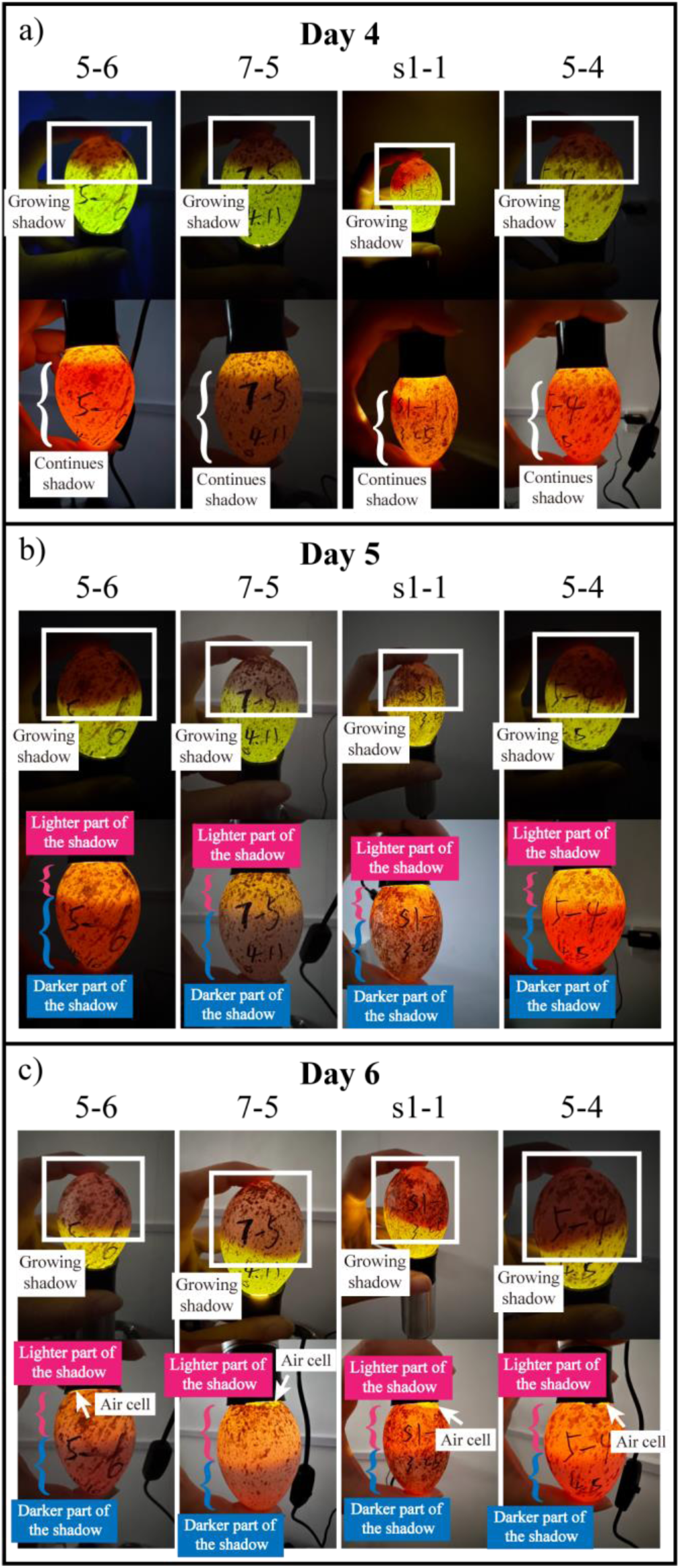
Representative candling images of fertile eggs during Day 4 and Day 6 of development. The growing shadows at the blunt end were highlighted in the white boxes. The white braces showed the continuous shadows at the sharp end. The darker and lighter parts of the shadow were pointed out by blue and pink braces separately. The white arrows showed the air cell at the blunt end. For each egg (5-6, 7-5, s1-1, 5-4), the images on the top were taken when candling from the sharp end, and those on the bottom were taken when candling from the blunt end.

**Day 7∼12:** The shadow at the blunt end when candled from the sharp end expands rapidly, occupying approximately half of the egg (Figure 3a-3f, white boxes). The position of the embryos may change if the incubation posture change (e.g. Change from vertical incubation with the blunt end upward to horizontal incubation in a flat position). Meanwhile, the air cell gradually enlarges and develops a distinct and well-defined boundary (Figure 3a-3f, white arrows).

**Figure 3.**
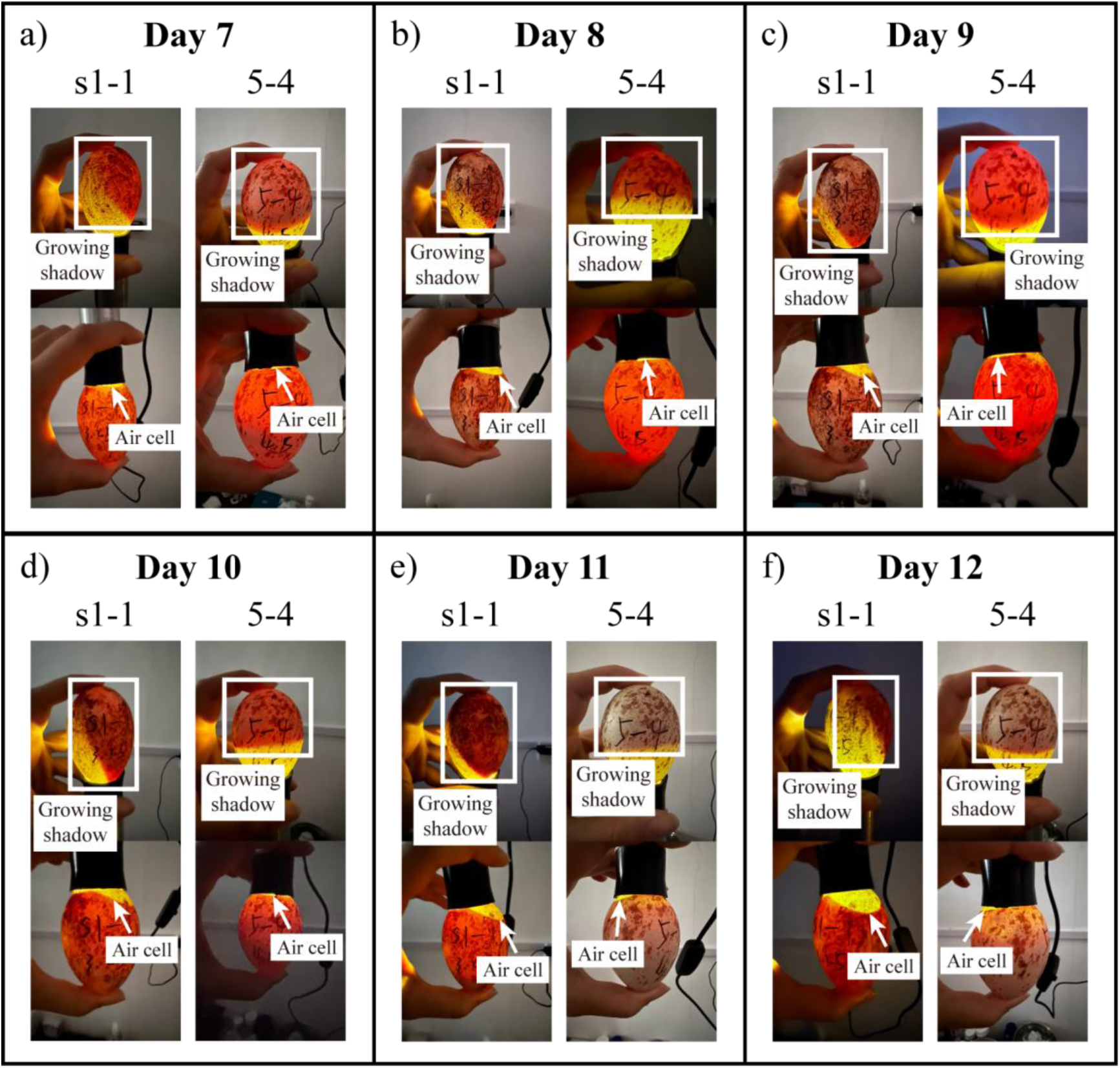
Representative candling images of fertile eggs during Day 7 and Day 12 of development. The shadows at the blunt end were highlighted in the white boxes. The air cells were pointed out by white arrows. For each egg (s1-1, 5-4), the images on the top were taken when candling from the sharp end, and those on the bottom were taken when candling from the blunt end.

**Day 13∼16:** Vascular structures become visible along the margins of the shadow when candled from either end of the egg (Figure 4a-4d, red arrows). The air cell remained stable and clearly visible (Figure 4a-4d, white arrows). When candled from the sharp end, occasional embryonic movements are observable (Figure 4a-4b, light blue arrows).

**Figure 4.**
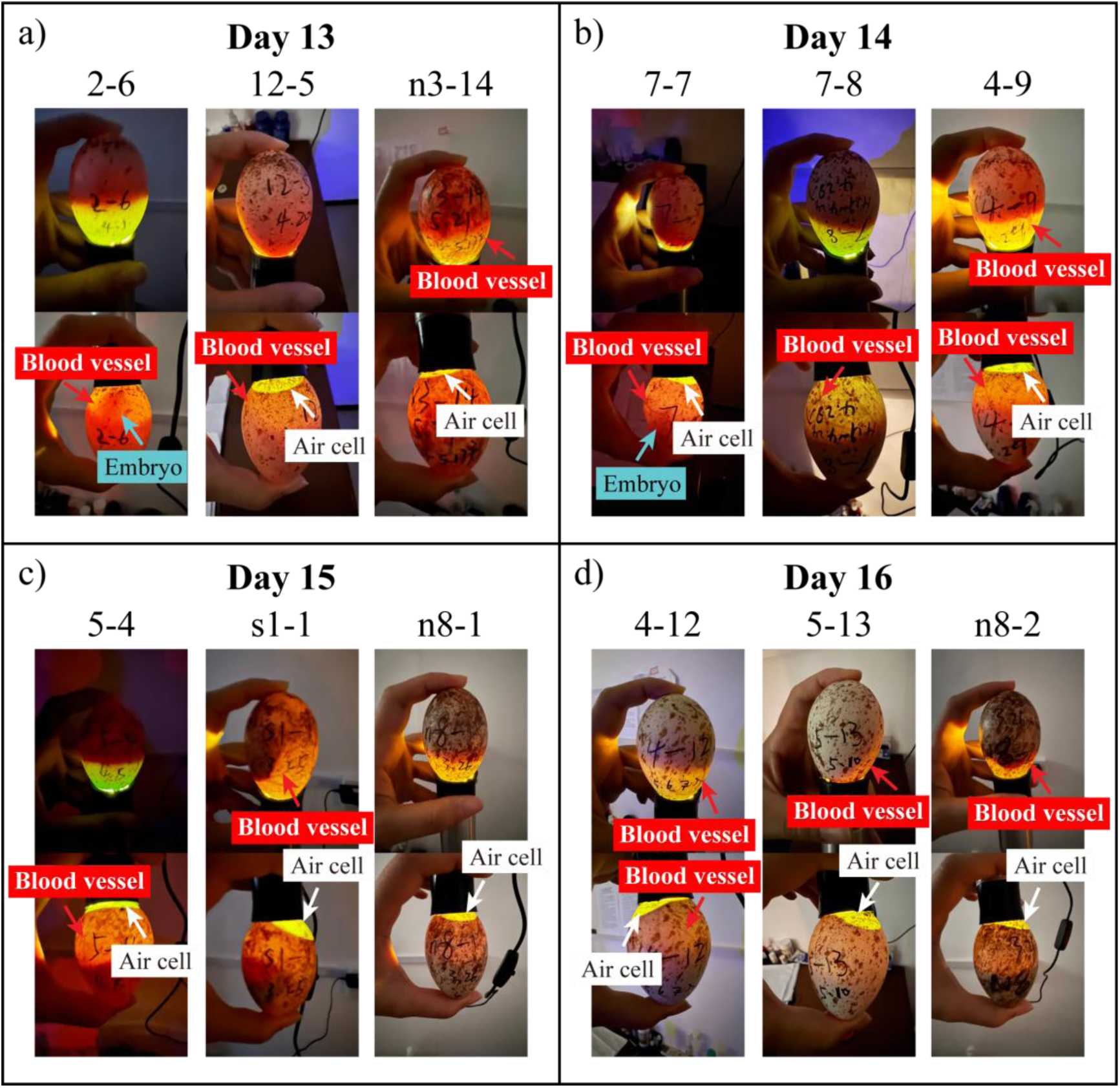
Representative candling images of fertile eggs during Day 13 and Day 16 of development. The red arrows showed vascular structures around the shadows. The white arrows pointed out the air cells. The embryonic movements were pointed out by light blue arrows. For each egg (2-6, 12-5, n3-14, 7-7, 7-8, 4-9, 5-4, s1-1, n8-1, 4-12, 5-13, n8-2), the images on the top were taken when candling from the sharp end, and those on the bottom were taken when candling from the blunt end.

**Day 17∼20:** The embryo shadow becomes opaquer, losing its reddish translucency (Figure 5a-5d, white boxes), and peripheral vasculature becomes more prominent (Figure 5a-5d, red arrows).

**Figure 5.**
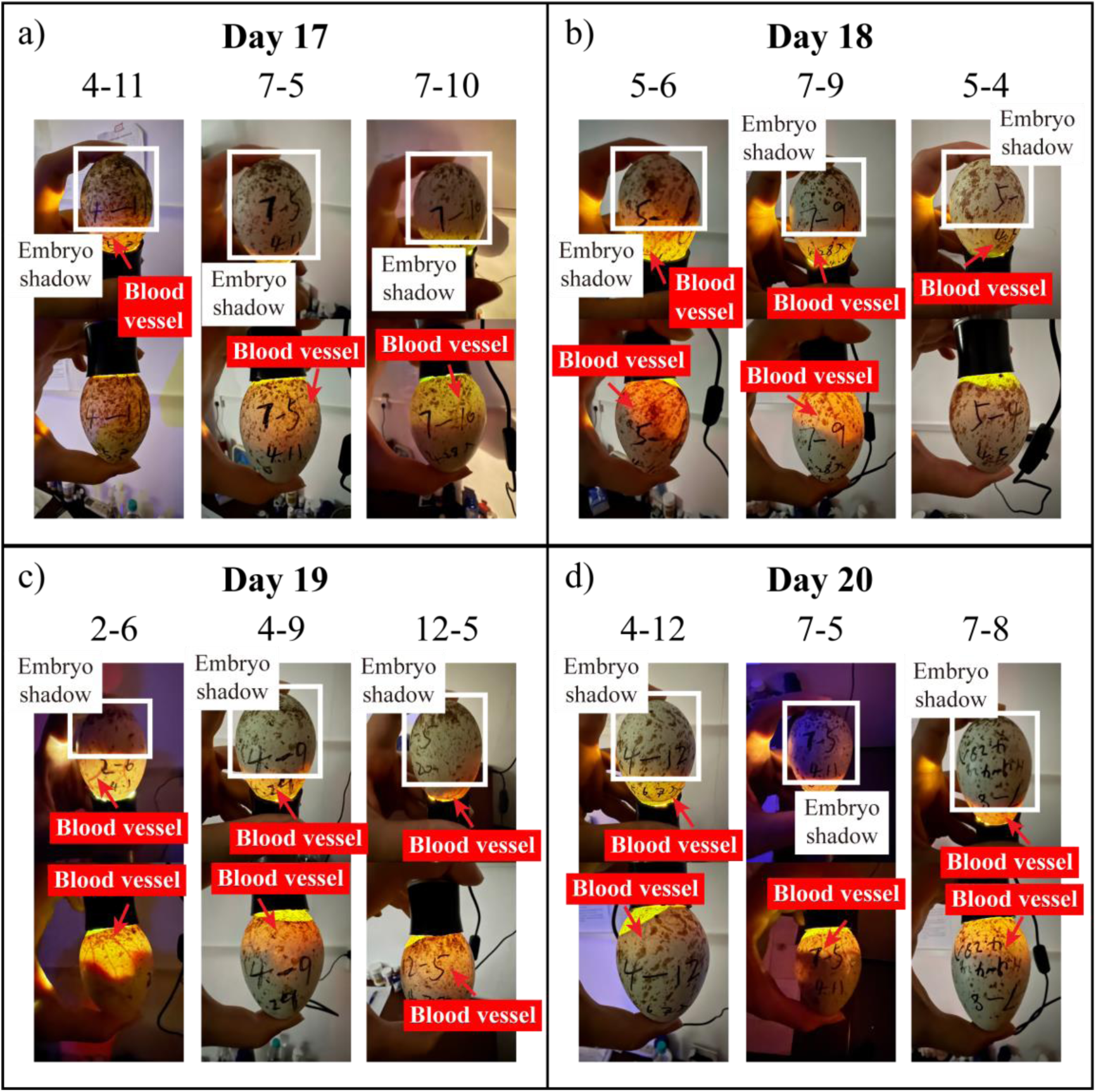
Representative candling images of fertile eggs during Day 17 and Day 20 of development. The white boxes showed embryo shadows. The red arrows pointed out the vascular structures. For each egg (4-11, 7-5, 7-10, 5-6, 7-9, 5-4, 2-6, 4-9, 12-5, 4-12, 7-8), the images on the top were taken when candling from the sharp end, and those on the bottom were taken when candling from the blunt end.

**Day 21∼24:** Vascular structures begin to regress as the embryo enlarged. When candled from the sharp end, the embryo shadow becomes nearly opaque, occupying almost the entire egg and greatly limiting light penetration (Figure 6a-6d, white boxes). When candled from the blunt end, the air cell continues to enlarged gradually with smooth and sharply defined boundaries (Figure 6a-6d, white arrows).

**Figure 6.**
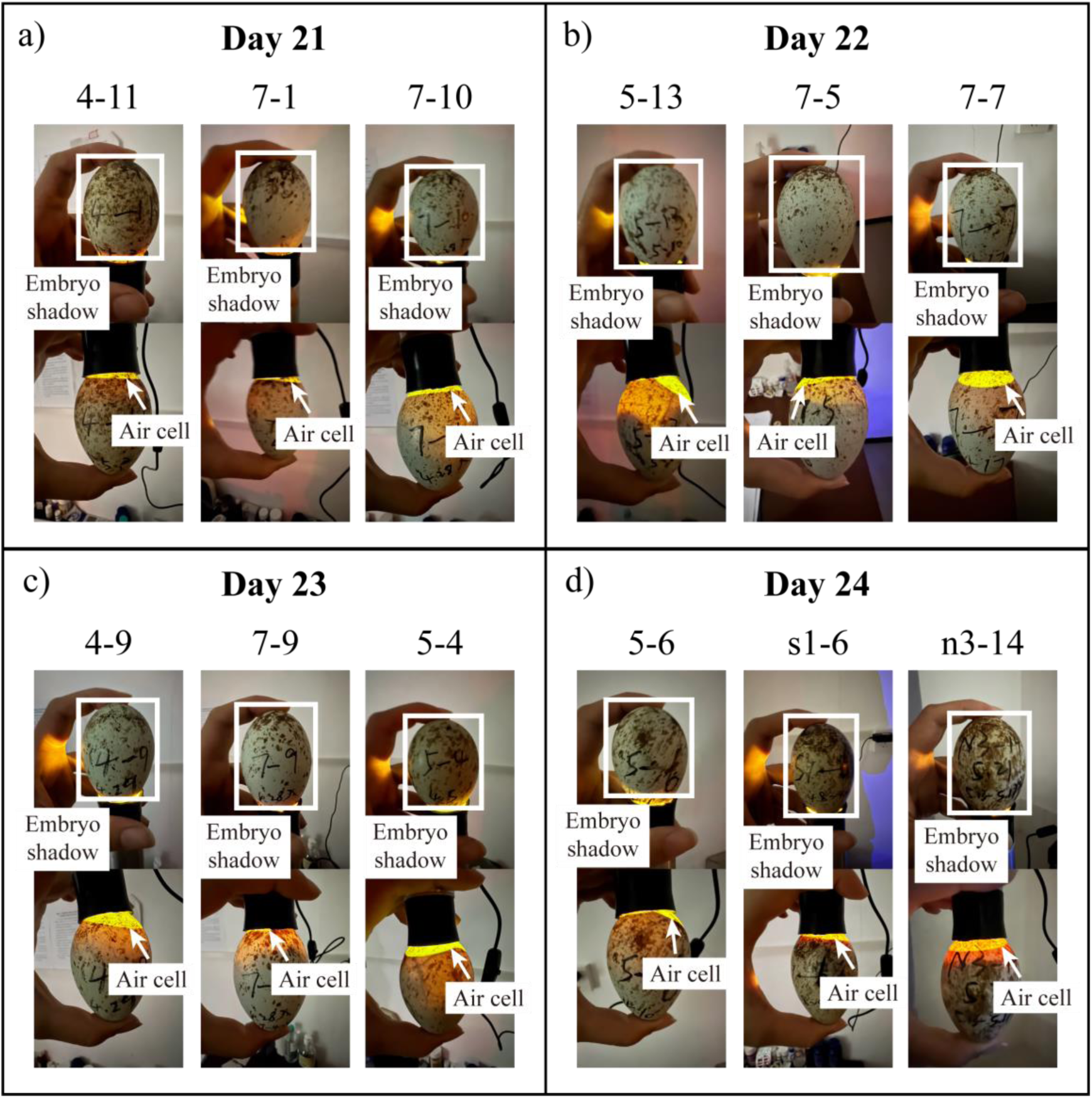
Representative candling images of fertile eggs during Day 21 and Day 24 of development. The embryo shadows were highlighted in white boxes. The air cells were pointed out by white arrows. For each egg (4-11, 7-1, 7-10, 5-13, 7-5, 7-7, 4-9, 7-9, 5-4, 5-6, s1-6, n3-14), the images on the top were taken when candling from the sharp end, and those on the bottom were taken when candling from the blunt end.

**Day 25∼29:** The embryo fills nearly the entire egg except for the air cell, which remains relatively stable in size with clear and well-defined margins. Pipping and hatching occur during this stage (Figure 7).

**Figure 7.**
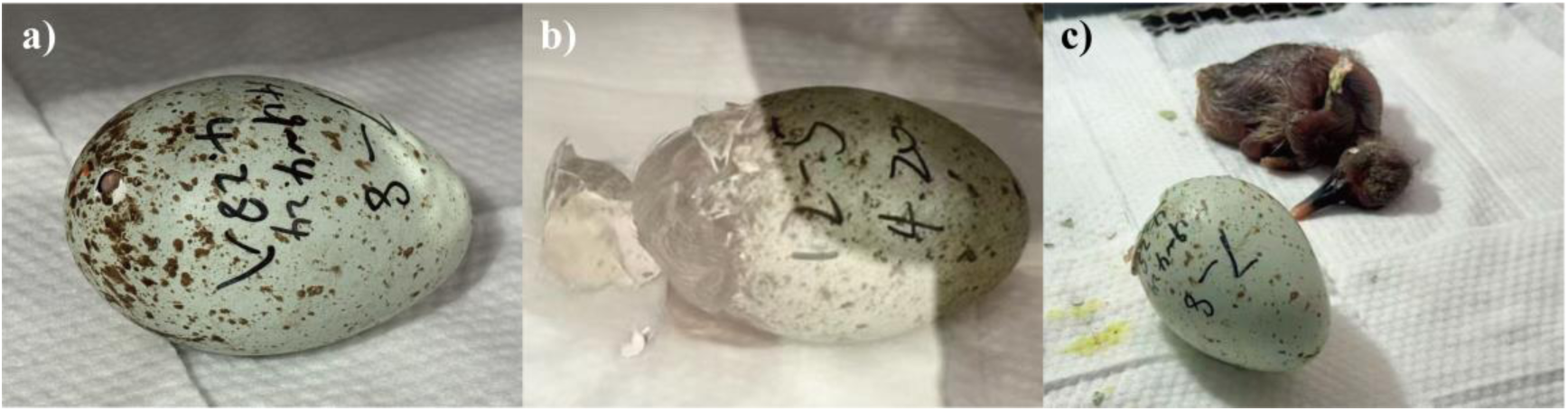
Hatching eggs and nestling that had just hatched out. a) Egg 7-8 on Day 26 when the bird started pipping; b) Egg 12-5 on Day 28 when the blunt end of the egg had been opened; c) Nestling 7-8 on Day 27 when it had just hatched out.

### b) identifying non-viable eggs

By comparing the candling images throughout incubation, we were able to distinguish viable and non-viable eggs (Figure S1), and remove the latter timely before they decayed and caused contamination. In the following sections, we present some representative images and provide the detailed descriptions that help differentiate between the various types of eggs.

#### fertile eggs vs. infertile eggs

With continuous candling observations, fertilized and unfertilized eggs can generally be distinguished during Day 4 to Day 6 of development (Figure 8a). When candled from the sharp end, the fertile eggs exhibit a progressively enlarging shadow caused by the developing embryo (Figure 8a, white boxes), whereas the shadow in the infertile eggs show little change (Figure 8a, red boxes). When candled from the blunt end, the shadow in the infertile eggs remains roughly spherical rather than occupying a defined area. In some cases, particularly in the later stage of incubation, the shadow in infertile eggs also appears to occupy a certain area (Figure 8b, red braces). However, these eggs lack clearly defined air cells with distinct edges, which are characteristic of fertile eggs (Figure 8b, white arrows), but only a vague cavity (Figure 8b, red arrows).

**Figure 8.**
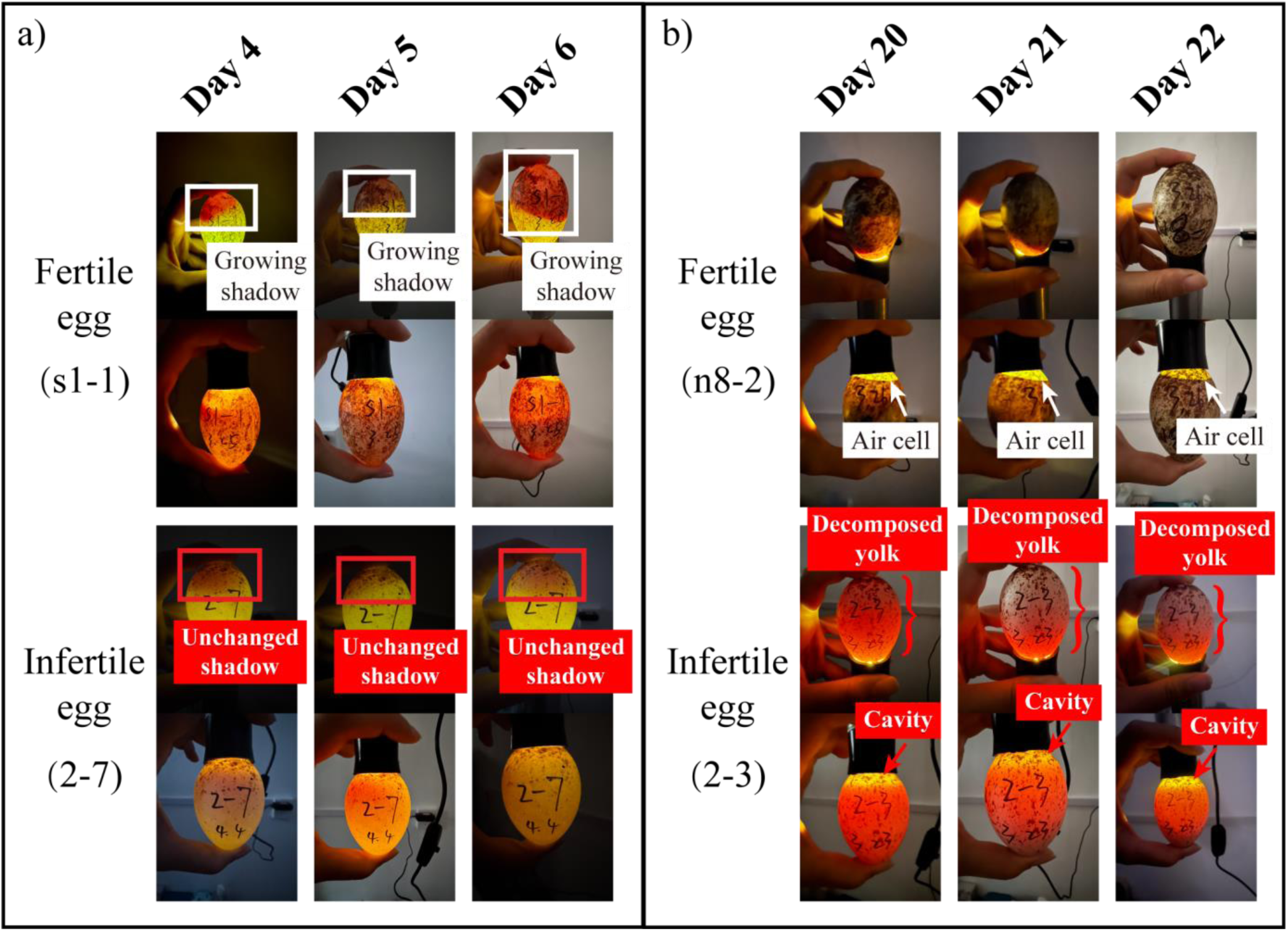
Representative candling images of fertile eggs (s1-1, n8-2) and infertile eggs (2-7, 2-3). a) Images taken during Day 4 and Day 6 of development, when the growing shadow of embryo (highlighted with white boxes) inside fertile egg became visible, while the infertile egg showed no significant changes (highlighted with red boxes); b) Images taken during Day 20 and Day 22 of development. In fertile eggs, the embryo’s shadow almost filled the entire egg except the air cell (pointed out with white arrow), whereas in infertile egg, the yolk had decomposed and formed a reddish shadow (pointed out with red braces). Meanwhile, the cavity (pointed out with red arrow) in blunt end lacks clear boundaries. For each egg (s1-1, 2-7, n8-2, 2-3), the images on the top were taken when candling from the sharp end, and those on the bottom were taken when candling from the blunt end.

#### embryonic shadow vs. decayed yolk shadow

When the eggs are not removed immediately after laying and continuous images of candling are unavailable, it may be difficult to distinguish fertilized from unfertilized eggs solely based on the presence of large shadowed areas. In fertilized eggs, the shadow gradually expands (Figure 9, white braces) as embryonic development progresses with a clear air cell remains at the blunt end (Figure 9, white arrows). During the mid-development stage, the embryo and its vascular network can be visualized under candling (Figure 4&5). In contrast, the extensive shadow observed in unfertilized eggs usually results from yolk decomposition and rupture, which mixes with the albumen. It typically appears abruptly within 1 or 2 days (Figure 9, images of egg 2-2 on Day 19 and Day 20), with indistinct edges, a relatively uniform appearance, and no observable structural features (Figure 9, red braces).

**Figure 9.**
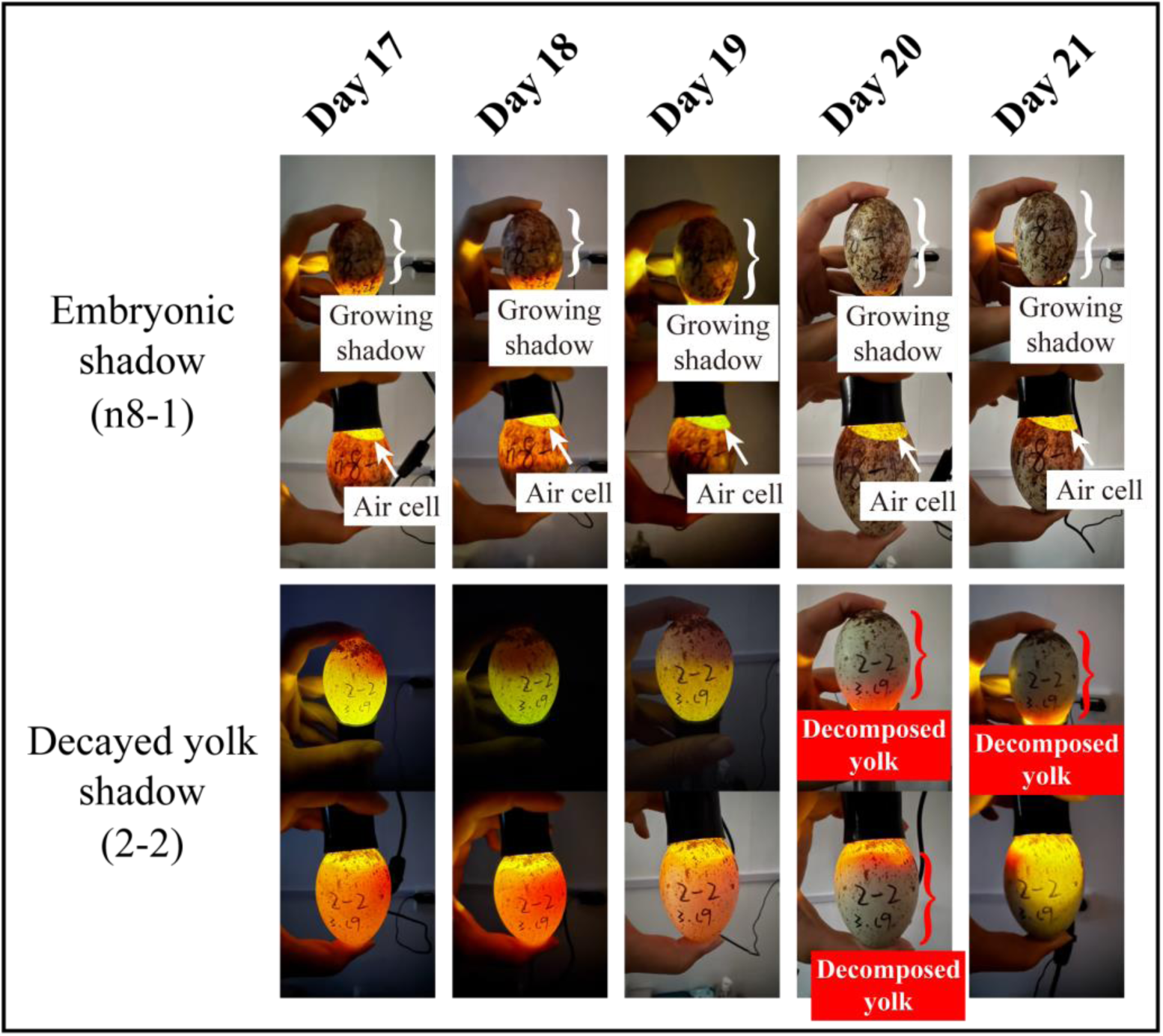
Representative candling images of fertile egg (n8-1) and infertile egg (2-2) during Day 17 to Day 21 of development. The growing shadow of embryo was highlighted with white braces and the air cell was pointed out by white arrow. The shadow of decomposed yolk without clear structure and boundaries was highlighted with red braces. For each egg (n8-1, 2-2), the images on the top were taken when candling from the sharp end, and those on the bottom were taken when candling from the blunt end.

#### living embryo vs. dead embryo

Apart from differences in growth, the shadows of living and dead embryos exhibit other distinctive features. Living embryo displays stable shadows and does not move even we shake the egg slightly. The edges of both the embryo shadow (Figure 10, white boxes) and the air cell are well-defined (Figure 10, white arrows), and occasional embryonic movements may be observed. In contrast, dead embryo shows shifting shadow, the edges of which display diffuse red structures resulting from ruptured blood vessels (Figure 10, red boxes), and the borders of the air cell are blurred (Figure 10, red arrows). When the egg is tilted, the fluid appears to invade the area of the air cell, indicating disrupted internal structures. As the vessels or yolk sac ruptures, the egg contents become turbid, and the candling image may resemble that of an infertile egg with a decomposed yolk (Figure 9&10, red braces).

**Figure 10.**
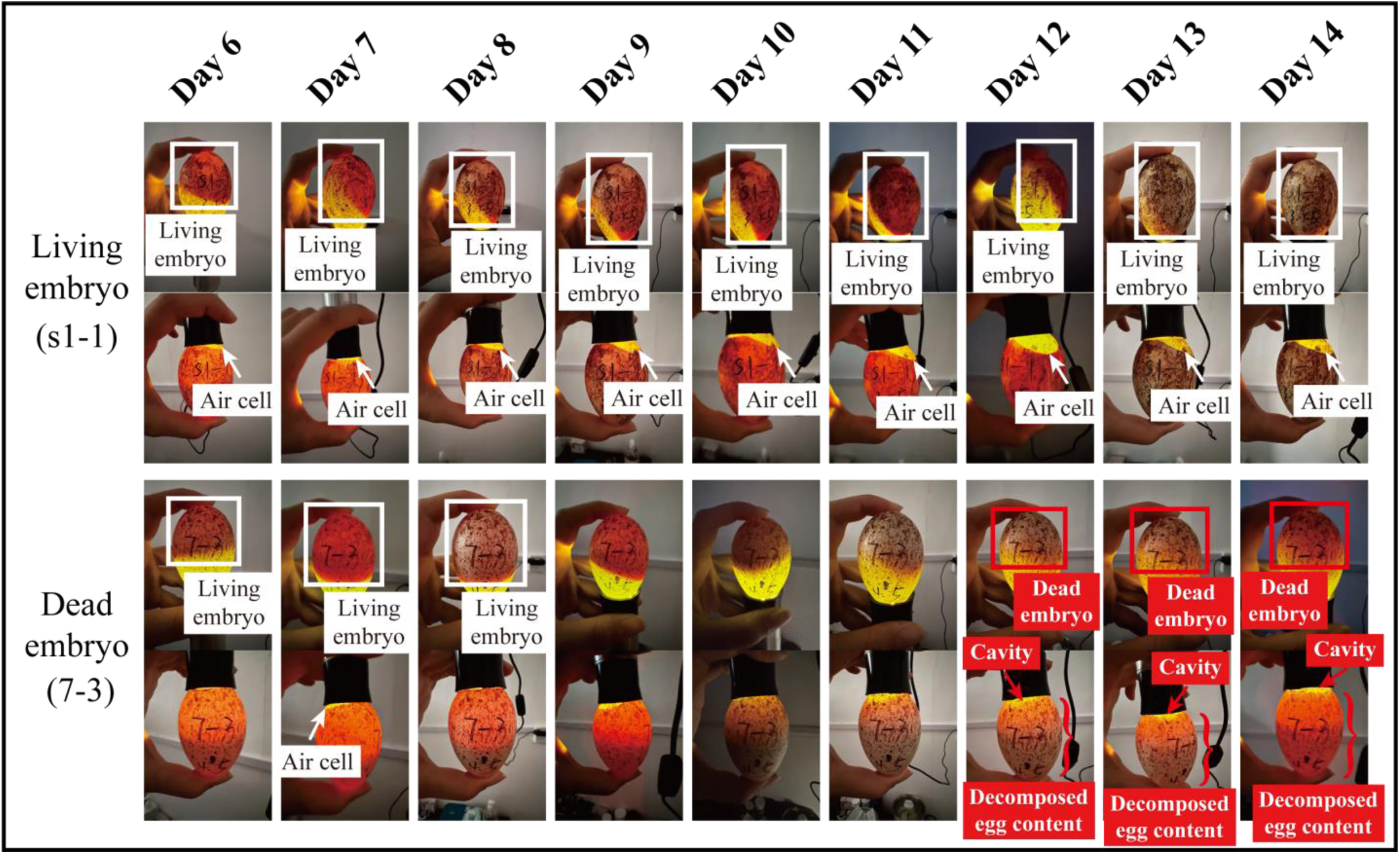
Representative candling images of living embryo (s1-1) and dead embryo (7-3, which died at around Day 10) during Day 6 to Day 14 of development. The living embryo was highlighted with white boxes, and the air cell was pointed out with white arrows. The dead embryo was highlighted with red boxes, and the shadow of decomposed egg content was pointed out with red braces. Meanwhile, the cavity (pointed out with red arrow) in blunt end lacks clear boundaries. For each egg (s1-1, 7-3), the images on the top were taken when candling from the sharp end, and those on the bottom were taken when candling from the blunt end.

### c) vital periods in embryo development

Based on four embryos with recorded death times (5-9, 4-6, n4-5, n3-11), developmental stages could be partly characterized as follows: Day 7, embryo was less than 2 cm, with distinct eye pigmentation, rudimentary limb buds, and visible heart chambers; by Day 9, the eyes had enlarged and the limbs were more prominent; by Day 15, the beak was clearly developed, the head was nearly equal in size to the body with large and prominent eyes, and the limbs were noticeably more advanced; by Day 23, feathers were already developed, the eyes were covered by eyelids, limb structures were fully visible, the beak had hardened, and egg teeth had appeared. Additionally, 3 embryos that died due to abnormal fetal position (n8-1, 5-4, 15-3) had their head and beak compressed or blocked by other body parts, preventing successful pipping (Figure S1).

Using these four reference embryos and the summarized differences in candling images between living and dead embryos, we estimated the death times for other 20 embryos (excluding 3 that died from accidents and 2 for which no egg-opening images were available) and placed them on the developmental timeline. Embryo mortality was found to be concentrated during mid-stage of incubation (Days 7∼15) (N = 12, 60.0%) and near hatching (Days 23∼29) (N = 7, 35.0%) (Figure 11), suggesting that these vital periods in the development of Crested ibis embryos requireparticular attention.

**Figure 11.**
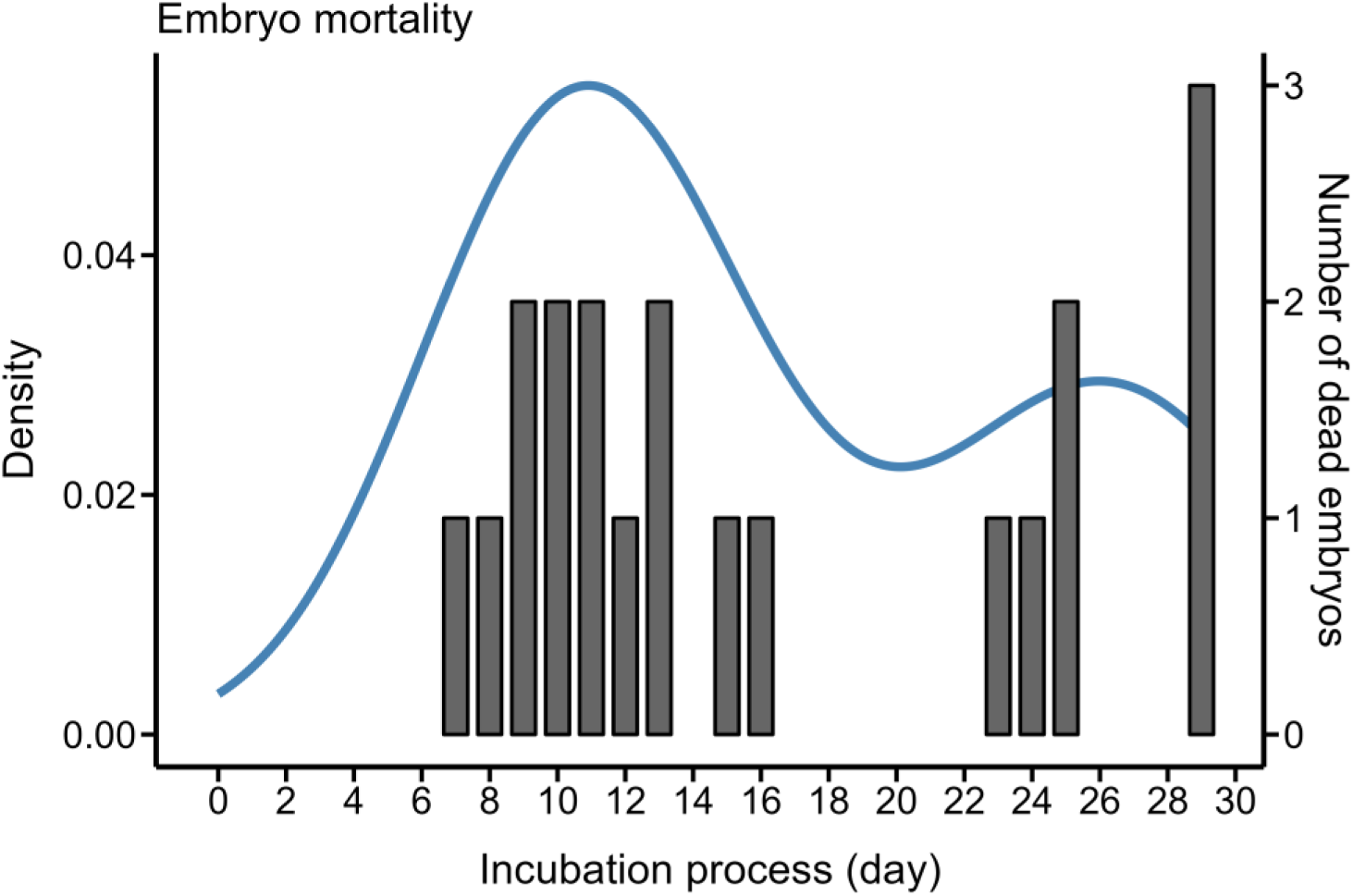
Estimated times of death for dead embryos (excluding 3 that died from accidents and 2 for which no egg-opening images were available). The left vertical axis represents the density distribution of dead embryos across different incubation ages, while the right vertical axis indicates the exact number of dead embryos recorded on each day.

## Discussion

In this study, we present the first comprehensive description of egg development, encompassing both the changes observed in unfertilized eggs and daily embryonic development in the endangered Crested ibis based on candling images throughout incubation (Figures 1∼7). By summarizing characteristic features at different developmental stages, we provide a practical reference for assessing fertilization status and embryo viability (Figures 8∼10), which can facilitate the early detection of abnormalities and timely management interventions. In addition, by integrating morphological characteristics with candling records, we estimated the timing of embryo mortality and found that most deaths occurred during mid-incubation (Day 7∼15) (N = 12, 60.0%) and shortly before hatching (Day 23∼29) (N = 7, 35.0%) (Figure 11, Figure S1). These findings reveal critical periods in the embryonic development of Crested ibis that deserve particular attention and provide an important basis for improving hatching success and understanding developmental constraints in this endangered species.

By comparing all the candling images, we found that, in general, the difference in shadow between infertile and fertile eggs became discernible after approximately 4–6 days of incubation (Figure 2). Based on the morphological information from other avian species with comparable incubation periods (∼28 days), the embryo at this stage shows a complete flexure from head to tail, with increasingly developed vascularization, and the early differentiation of major organs and body structures including the eyes, heart, and limbs (Fant 1957, Caldwell & Snart 1974, Araújo *et al*. 2019, Pisenti *et al*. 2024). The actual onset of embryonic development should occur earlier, but it can only be directly confirmed through a breakout examination. Between Day 7 and 12, the embryonic shadow gradually enlarged and appeared as a relatively transparent reddish area during candling (Figure 3). Based on the egg-opening images of embryos that died unexpectedly and developmental patterns reported in other studies (Caldwell & Snart 1974, Araújo *et al*. 2019, Pisenti *et al*. 2024), it is known that the embryo remains relatively small at this stage, with a developing but not yet dense vascular network. Consequently, the candling images display a relatively transparent shadow with a distinct reddish hue. From Days 13 to 16, clearly defined vascular structures became visible at the periphery of the embryonic shadow (Figure 4). The shadow gradually darkened, and in some cases, slight embryonic movement could be observed. However, due to the natural coloration of Crested ibis eggs, which are covered with dark speckles and have relatively low shell translucency, it was difficult to distinguish specific structures within the shadow. On Day 17 to 20, the embryonic shadow expanded further, the vascular network became more distinct, and embryonic movements were more easily detected (Figure 5). By Days 21 to 24, the shadow had turned almost completely opaque, occupying nearly the entire egg (Figure 6). At this stage, the embryo had developed distinct and clearly visible feathers, while more translucent structures such as blood vessels and yolk sacs were gradually absorbed or disappeared (Figure S1). In species with similar developmental processes, embryos at this stage are also increasingly well-developed. Feathers grow rapidly, scales gradually become complete, and the yolk sac is progressively withdrawn into the body cavity (Fant 1957, Caldwell & Snart 1974, Araújo *et al*. 2019, Pisenti *et al*. 2024). Between Days 25 and 29, the embryo was fully developed. The chick inside began to chirp and peck at the shell. The time from the first pip to complete hatching could vary from 1 to 3 days.

Regarding the features previously reported in studies on chickens that can be used to identify non-viable eggs, the typical indicator of early embryonic death is the appearance of a “blood ring” when inspecting the eggs (Ernst *et al*. 2004). This disorder has been described as an autosomal recessive lethal trait (Savage *et al*. 1988). In our study, we did not observe such a distinct “blood ring” when candling. This may be attributed to the relatively dark coloration and low translucency of the eggshell, as well as the presence of dark spots on the surface, which may hinder observation. Interspecific differences in the manifestation of embryo death might also account for this discrepancy. When embryos die after a period of development, the major indicators of death are typically a large, dark, round shadow and the absence of visible blood vessels (Ernst *et al*. 2004). This pattern is consistent with our observations in Crested ibis eggs that experienced embryonic death, where vascular structures disappeared and the edges of the shadow became blurred after death (Figure 10). Overall, most existing guidelines for identifying non-viable eggs through candling have been developed primarily for economically important species such as chickens and turkeys, underscoring the need for further research on a wider range of species, especially those that are protected.

Based on the egg-opening images of four embryos with recorded death times, we estimated the timing of death for most embryos by comparing their images taken during incubation and after egg opening. Our results indicated that embryo mortality primarily occurred during the mid-incubation stage (Day 7∼15) and near hatching (Day 23∼29). Previous studies have reported that avian embryos are more likely to die during the early and late stages of incubation (Romanoff 1949). Research on a threatened bird, the Hihi (*Notiomystis cincta*) revealed that early embryo mortality (Day 0∼5) accounted for an average of 56.8% (±15% SD) of total embryo deaths (Morland *et al*. 2024). In the study conducted by Yang et al. on the artificial breeding of Crested ibis in Sichuan (2017 ∼ 2019), the highest mortality occurred during Day 17∼25 (57.7%–75.8%), followed by Day 9∼16 (9.1%–34.7%) (Yang *et al*. 2020). The differences in the conclusions of different studies may be related to interspecies differences, the different division of the incubation stage, and the differences in the incubation environment. Common non-genetic causes of embryo death include nutrient deficiencies in the eggs, bacterial contamination, abrupt changes in temperature or humidity, inadequate or improper turning, and so on (Wilson 1997, Ori 2011, Rideout 2012). In the current artificial breeding process of Crested ibis, procedures such as manual egg collection and cooling, as well as micro-environmental differences among incubators, could contribute to these mortality factors. Therefore, identifying the specific causes of hatching failure based on assessments of egg condition and developmental stage would facilitate targeted improvements in management practices. Additionally, we noticed that three embryos (Figure S1, n8-2, 5-4 and 15-3) that had fully developed and exhibited pre-hatching behaviors, such as chirping and shaking, ultimately failed to hatch and died. Dissection revealed that their heads and beaks were obstructed by other body parts, preventing successful pipping (Figure S1). During the late stage of incubation, rotate to position themselves for hatching, and factors such as rough handling and improper turning can disrupt this process (Rideout 2012). Small egg size may also be a contributing factor, as limited internal space can restrict the embryo’s movement. In our observations, egg 15-3, which was produced by a pair of one-year-old birds, was notably small, measuring only 62.92 mm in length and 40.78 mm in width—both shorter and narrower than the average dimensions of normally hatched eggs (66.59 [62.4∼71.28] mm and 45.92 [41.3∼48.5] mm). Reported length and width in previous studies were approximately 67.55 mm and 45.24 mm (Shi & Yu 1989, Xi *et al*. 2001). The confined space likely restricted the embryo’s movement, hindering the rotational behavior necessary for pipping.

Although the current population of the Crested ibis has greatly increased, the species still faces severe inbreeding and low genetic diversity, as the entire existing population originated from only seven individuals and two breeding pairs. Previous research showed that the nucleotide diversity of existing populations of the Crested ibis is markedly lower than that of the historic population, a substantial accumulation of deleterious mutations has occurred over the past four decades, posing potential genetic risks (Feng *et al*. 2019). Additionally, inbreeding in Crested ibis has already been found to be associated with increased embryonic mortality, and several candidate genes linked to detrimental diseases were identified in genomic regions that differ between viable and dead embryo samples (Fu *et al*. 2019), potentially compromising embryo quality and increasing susceptibility to environmental fluctuations during incubation. Therefore, it is essential to accurately determine the fertilization status of eggs and the survival status of embryos in order to obtain reliable estimates of fertilization and embryo survival rates, which are critical for elucidating how and to what extent inbreeding affects the reproductive success of the Crested ibis. At the same time, these can also be compared and discussed in relation to those reported in other species.

At present, studies on the incubation and embryonic development of Crested ibis remain relatively limited. Most previous research has focused primarily on quantitative metrics such as clutch size, fertilization rate, and hatching rate, but has lacked detailed observations and descriptions of the embryonic development process (Liu *et al*. 1999, Xi *et al*. 2001, Huang *et al*. 2006, 2016). Although candling has been applied in practical artificial breeding, the absence of standardized descriptions for candling results means that assessments of egg fertilization status and embryonic development often rely heavily on the experience and skill of individual staff, lacking uniform criteria. Without systematic references, the risk of human error might increase. Furthermore, to avoid prematurely terminating the incubation of fertilized eggs, all eggs are often incubated for extended periods, which may result in dead embryos or infertile eggs deteriorating and contaminating the incubator if not removed in a timely manner. While developmental patterns from closely related species can provide some guidance, they are less effective than species-specific information. Based on 1,423 images from 106 eggs, we have preliminarily summarized the egg development patterns of Crested ibis, providing detailed descriptions and illustrative references. This resource can aid researchers and breeding staff in accurately aging normal embryos and promptly identifying and removing abnormal eggs, thereby reducing contamination risks in artificial incubators.

Strictly speaking, a complete description of egg development should also include detailed morphological data obtained through direct examination of fertilized eggs, as has been conducted in previous studies on other avian species (Sellier *et al*. 2006, Nagai *et al*. 2011, Hemmings & Birkhead 2016). However, due to conservation constraints, it is currently not feasible to acquire such detailed morphological data for the Crested ibis through invasive methods like egg dissection. Meanwhile, in our study, the examination of infertile eggs and those containing dead embryos was conducted only after confirming that there were no signs of life. By that time, most eggs had already begun to decay, and many embryonic structures were no longer clearly distinguishable, which inevitably limited the accuracy of our descriptions. This may also have resulted in some fertilized eggs that died at very early stages (Day 0∼3) being misclassified as unfertilized eggs due to the lack of visible embryonic features in the candling images, thereby leading to a possible overestimation of the infertility rate and an underestimation of early embryonic mortality. Therefore, continued accumulation of data and non-invasive observation methods will be essential to refine our understanding of Crested ibis egg development. Only through detailed documentation of both normal developmental stages and the characteristics of embryos in different conditions can we gain a comprehensive understanding of developmental patterns, apply this knowledge to practical management, and enable comparative analyses across species.

## Conclusion

We present the first description of egg development in Crested ibis based on candling images throughout incubation. Our study also provides a practical framework for assessing fertilization status and embryo viability, including indicators such as the growth rate and shape of the shadow, and the presence of clearly defined air chambers, which can facilitate the early detection of abnormalities and enable timely management interventions. Furthermore, by integrating morphological characteristics with candling records, we estimated the timing of embryo mortality and found that most deaths occurred during mid-incubation (Day 7∼15) and near hatching (Day 23∼29), which reveals the critical periods of embryonic development that warrant particular attention. Overall, our findings offer an important basis for improving hatching success and enhancing understanding of embryo developmental constraints in this endangered species.

## Notes

### Competing Interest Statement

The authors have declared no competing interest.

### Funding

- Institute of Zoology, Chinese Academy of Sciences: 2023IOZ0104, 2024IOZ0107
- National Natural Science Foundation of China: General Program, Grant No. 32570596

