## Supplemental Figure 1 for "Candling analysis of egg development in an endangered bird species Crested ibis (*Nipponia nippon*)"

Figure S1. The dead embryos of the Dongzhai Crested ibis population in 2025 breeding season with images after break-out examination. Egg 5-9, 4-6, n4-5 and n3-11 died in accidents with documented death times. Egg n8-1, 5-4 and 15-3 died from abnormal fetal position. The times of death for other embryos were estimated based on their candling images and the morphological features.

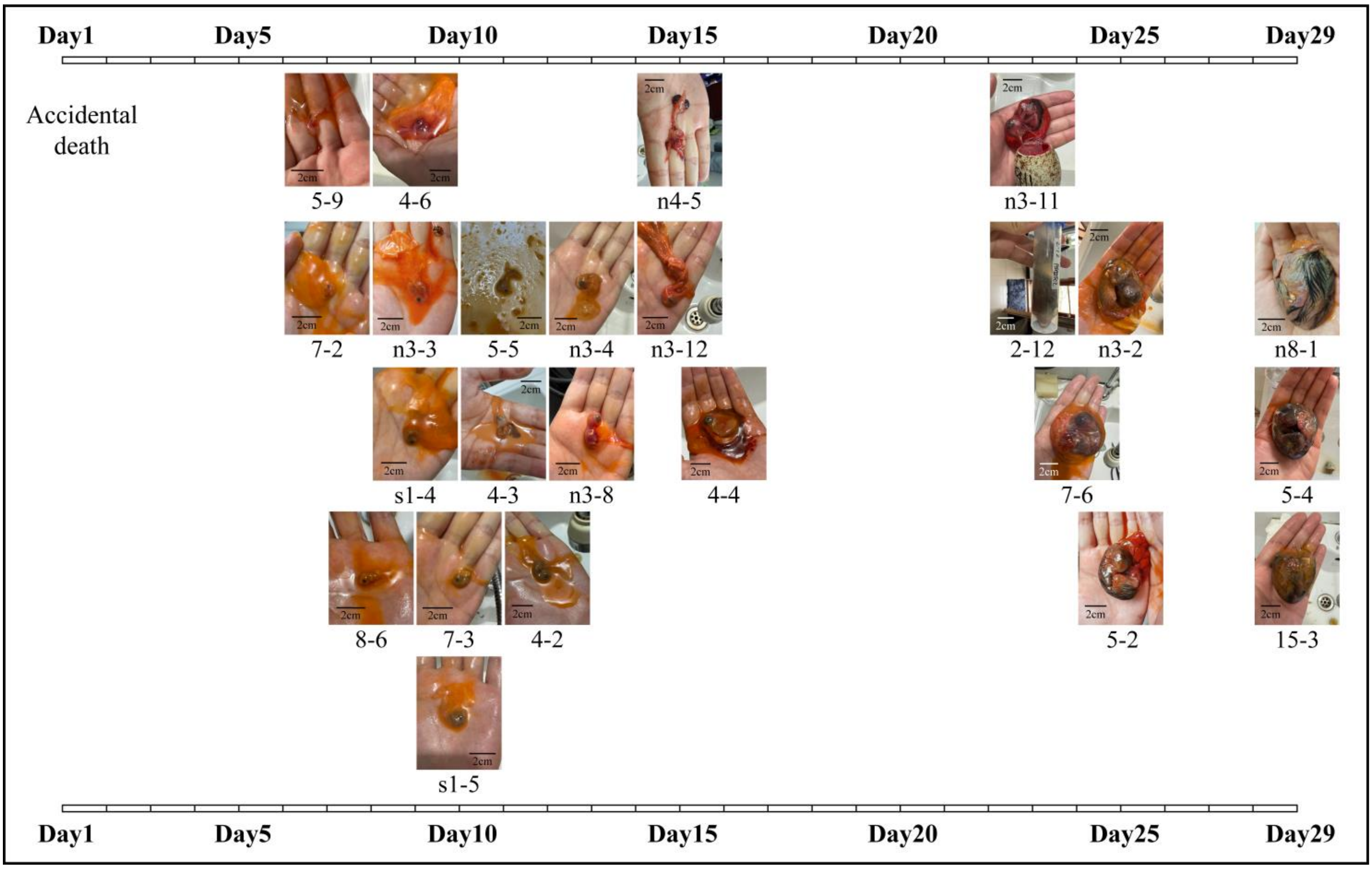
